# SWIF-TE: identifying novel transposable element insertions from short read data

**DOI:** 10.1101/2025.07.30.667279

**Authors:** Claire C Menard, Nathan S Catlin, Adrian E Platts, Yinjie Qiu, Emma Roback, Manisha Munasinghe, Nathan M Springer, Emily B Josephs, Candice N Hirsch

## Abstract

Transposable element (TE) insertion polymorphisms (TIPs) are TEs not in the same location between individuals. TIPs have contributed to genomic and phenomic variation but have been historically difficult to study due to their repetitive nature. Here, we describe a fast and memory-efficient tool to identify novel TIPs from short read sequences. SWIF-TE was able to identify 1,438 insertions at a precision rate of 27% using 0.10 Gb of memory and 0.82 hours of runtime from 15x resequencing data of a non-reference maize inbred. SWIF-TE is a powerful tool for studying TE variation in species with TE-rich genomes.

## BACKGROUND

Transposable elements (TEs) are DNA sequences that can move around within a genome, and they have important implications for genome evolution. TE insertions can cause changes at the sequence level (Xia et al., 2024), alter transcription factor binding (Merenciano & González, 2023), and affect methylation patterns (Noshay et al., 2019). Variation caused by TE insertions has been demonstrated to have phenotypic consequences across species (Wells & Feschotte, 2020). For instance, the insertion of a *Rider* TE within the promoter region of a flowering time gene in *Arabidopsis* contributed a new TF binding site that altered gene regulation (Quadrana et al., 2019) and resulted in later flowering time. A gene responsible for tomato flavor harbors a TE insertion that was present in the first cultivar released in Europe in the 16th century and was not found in wild relatives (Domínguez et al., 2020). In soybean plants, the excision of a *CACTA-* like TE resulted in flowers with different colored sections (Xu et al., 2010). Likewise, the peppered moth phenotype that occurred during industrialization in Great Britain was the result of a large tandemly repeated TE insertion in the *cortex* gene (Van’t Hof et al., 2016). Despite the significant role of TEs in the genomes of many species and the potential effects of TE insertional polymorphisms (TIPs) on phenotypic variation, there is still limited knowledge about population-scale within-species variation due to the computational challenges of studying these highly repetitive parts of the genome.

Recent advancements in methodology have significantly improved our ability to assess TE insertions at a whole-genome level. The most accurate approach is to utilize whole-genome sequence aligners alongside available TE annotations to identify novel insertions and deletions between pairs of individuals. Such techniques have been applied to multiple genome assemblies in maize revealing the dynamics of TE behavior within the species (Munasinghe et al., 2023). While whole-genome alignment methods are highly effective, they are prohibitively expensive for population-scale studies in species with large genomes. With the advancement of long read sequencing technology, new programs have been developed to identify structural variants (SV) and TE insertions between genomes <u>(Groza et al. 2024; Mohamed et al. 2023)</u>. Long read programs often rely on a single read containing the entire TE sequences as well as the surrounding insertion site sequence. While these approaches are quite promising, the acquisition of sufficient samples, extraction of high molecular weight DNA, and the cost of generating long-read data still limit the application in large populations of individuals.

Short read data is currently the most efficient and cost-effective method available to study TIPs on a genome-wide population level. Many approaches have been developed using short reads that can be used to either score the presence/absence of known (annotated) TEs from a reference genome in a population of individuals (Cai et al., 2022; Qiu et al., 2021) or to identify the position of novel insertions not present in the reference genome assembly (Baduel et al., 2021; Sohrab et al., 2021; Yu et al., 2021). Identifying novel insertions is more difficult because the sequence is not present in the reference during mapping. Still, short read data can provide evidence of these insertions. There are generally three different types of evidence used from short read sequence mapping to identify coordinates of novel TE insertions:split, discordant, and anchored reads. Multiple programs have been developed to identify coordinates using these different approaches, though with various limitations <u>(Orozco-Arias et al. 2020;</u> <u>Sohrab et al. 2021; Yu et al. 2021)</u>.

Anchored paired-read data looks for instances in which one read aligns to the genome and the other aligns to the TE library. A notable program that uses this approach is TIP_finder (Carpentier et al., 2019). This tool was developed for use with the rice genome, which is approximately 0.43 Gb with 46% TE content (Li et al., 2024). Application of this program to 3000 rice genomes identified over 50,000 TIPs that were used to identify the origin of rice domestication from three distinct events. While this approach works for smaller genomes, it becomes less effective upon scaling up to larger genomes.

Other tools have been developed using split and discordant read data to identify TIPs. Split reads are reads that do not map contiguously to the reference genome, but rather have two partial alignments. Discordant read pairs are those that both map fully to the genome, but whose alignment to the reference have a different orientation or distance as compared to the reference. The TEMP2 pipeline (Yu et al., 2021), hosted by the McClintock package ((Nelson et al., 2017) uses split and discordant reads and clusters them into fragment sizes to identify novel TE insertion sites. Conversely, TEfinder (Sohrab et al., 2021) uses just discordant reads to identify novel TE insertions. TEMP2 was first used to find insertions of the *HERV-K* retrovirus family that are associated with human cancer (Orozco-Arias et al., 2020), and TEfinder was used to find small *Hornet* transposon insertion events in *Fusarium oxysporum* (Sohrab et al., 2021). Both of these tools were originally developed to run on one TE family at a time and were not designed for genome-wide analyses. Other tools, such as SPLITREADER2.0 (Baduel et al., 2021) and MELT (Gardner et al., 2017), were designed to use both discordant and split-reads to identify new TE insertions and deletions of all TE families. SPLITREADER2.0 uses the SPLITREADER (Domínguez et al., 2020) script to identify TE insertions and TEPID (Stuart et al., 2016) to identify TE deletions using discordant reads. It starts by mapping soft clipped split reads and discordant reads to a known TE library, remaps those reads to the reference genome, and then identifies insertions or deletions using target site duplication information. This program has been implemented at a population scale in tomatoes and found TE insertions responsible for favorable flavors in the process of domestication (Domínguez et al., 2020). MELT uses discordant read pairs to identify the TE family and split reads to get the target site duplication location (Gardner et al., 2017). It was implemented in a population-level human study that found recently mobile *FL-L1* elements in human cancer cells (Gardner et al., 2017). While these tools identify TE insertions from all TE families, they were not built to efficiently run on large populations of individuals in species with large genomes.

Thus, while there are a number of existing approaches and tools for analyzing TIPs with short read resequencing data, they were designed for use in small genomes with relatively low levels of TE content or to only identify TIPs for a limited number of TE families. There is no available program designed to handle the demands of genome-wide searches of all TE families in large genomes with high levels of TE content. As such, we created a new tool, SWIF-TE (Short Read Whole Genome Insertion Finder for Transposable Elements), which is optimized for use in large genomes, with a high percentage of TE content in the genome, that can be implemented at the population level. We tested SWIF-TE in maize, a species with a large genome, a high percentage of TE content, a high prevalence of polymorphic TEs, and with access to many high-quality genome assemblies and known TE insertions available to provide ground truth data. We investigated precision, false discovery rate (FDR), sensitivity, and F1 score in SWIF-TE and compared them to several other tools that were benchmarked. We also assessed the extent to which information across individuals in a population can be used to improve the discovery of novel TE insertions. We show that SWIF-TE is a computationally efficient program that is optimized for identifying novel TE insertions at the population level in species with large TE-rich genomes.

## RESULTS

### SWIF-TE is a tool to identify TE insertion polymorphisms using short read data

SWIF-TE is a whole-genome TE insertion finder that uses short read resequencing data as the input (Figure 1). The program, which is written in Xojo (https://xojo.com/), uses short reads aligned to both a reference genome assembly and a reference TE library. It sequentially loads in blocks of this sequence information, and identifies reads that have one end mapped to the reference genome and one end mapped to the TE library using the split-read approach. The output includes both the insertion point in the genome and the TE family for the putative insertion. SWIF-TE is fast and memory efficient, enabling it to be used on large, complex genomes at the population level.

**Figure 1.**
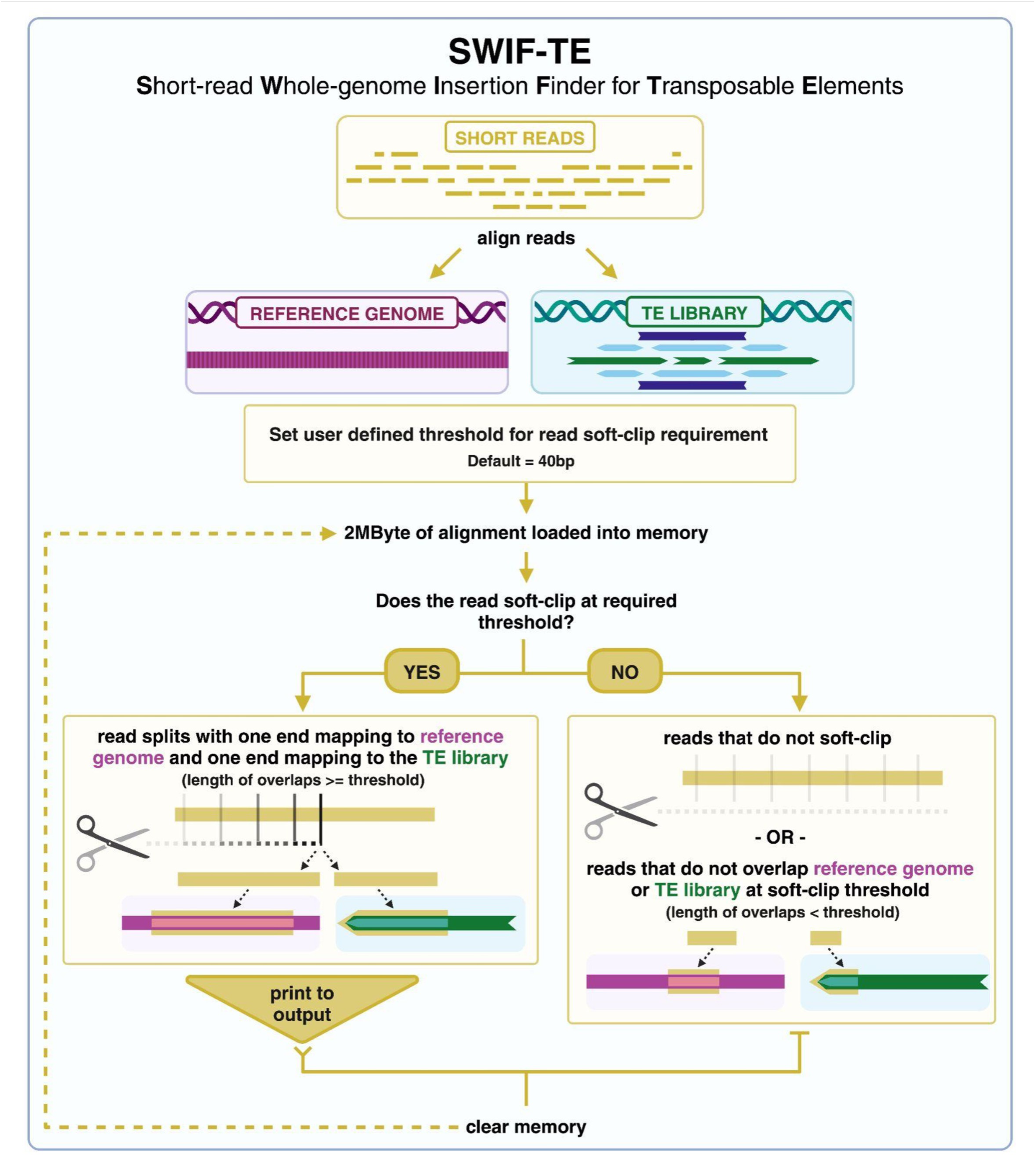
Schematic of the SWIF-TE algorithm. SWIF-TE begins by processing 2MB of alignment data from short reads aligned to the reference genome assembly and the TE library. A user-defined threshold for soft-clipping is set, and the alignment is processed through this filter. If one end of the read aligns to the reference genome assembly and the other end aligns to the TE library, then the alignment breakpoint (1bp) in the reference genome and the name from the TE library are printed to the output. If the alignment does not meet the user-defined threshold for soft clipping, or the reads do not align to the reference genome and the TE library, then the memory is cleared, and no information is printed to the output.

SWIF-TE requires two sequence alignment map (SAM) alignment files generated by aligning cleaned (i.e., quality and adapter trimmed) short reads (ideally 150bp or longer) to 1) a reference genome assembly and 2) a species-specific TE library using the bowtie2 aligner (Langmead & Salzberg, 2012). The TE library is a database of canonical TE sequences for the species or a related species. Annotation of the sequence in the database with the TE family name is used for downstream annotation of the putative TE insertion. Before running SWIF-TE, it is important to make sure there are no PCR duplicates in the input SAM file, as these duplicate reads could inflate the number of false positives in the output of SWIF-TE.

The SWIF-TE algorithm processes 2Mb blocks of alignment data at a time and breaks these blocks down into individual reads that are processed through an increasingly stringent set of criteria in order to reject uninformative reads as rapidly as possible. An informative read is one that maps uniquely at one end to the reference genome, while the other end is soft-clipped by the aligner. The algorithm then looks for a complementary alignment pattern in an alignment to the TE library, where the end that did not map to the reference instead maps into the TE library, and the other end is soft-clipped. Depending on the species of interest and the depth of sequencing, processing the SAM file can take substantial runtime and memory. However, SWIF-TE can be run in parallel by chunking the SAM file into 2 Mb blocks. The output for informative reads is a reference genome position where the read alignment ends, suggesting the insertion of a non-reference sequence at this point. The TE sequence name from the TE library and the length of the alignment to the TE sequence are also reported. The length of the alignment to the TE sequences provides a metric to assess the likelihood of the TE identity being correct and is one of many built-in filtering parameters described below.

Within the algorithm, there are many potential parameters that can be optimized. Alignment heuristics such as minimum MAPQ scores are hard-coded (minimum MAPQ score of 4) into the program, while other parameters that may be more species-dependent can be adjusted by the user to optimize for the species of interest. For example, SWIF-TE allows for custom alignment length criteria for alignments to both the genome assembly and the TE library. The default minimum alignment lengths are 40bp into the genome and 40bp into the TE library. When aligning longer reads (e.g., 200-250bp reads) or aligning to a species where TE content is dominated by a few families with highly similar termini, it may be helpful to increase the required alignment length. After running SWIF-TE, there are a number of options that can be applied for downstream processing of the informative reads. For example, sets of reads that report insertions at similar locations can be clustered within user-defined distances (e.g., within 50bp of each other) to concatenate the available evidence for an insertion. The number of reads supporting an insertion can also be optimized based on accuracy needs for downstream applications. Sequencing depth and species of interest could affect the level of read support needed to call an insertion precisely.

### SWIF-TE performance and optimization

To assess the performance of SWIF-TE, we sought to use a species with a TE-rich genome and substantial genomic resources to generate a gold-standard set of TE insertions to compare with SWIF-TE results. The maize genome is composed of 88% TEs (Ou et al., 2024), and there are multiple reference-quality genome assemblies with high-quality TE annotations (Hufford et al., 2021; Ou et al., 2024). This, coupled with the rich history of TE research in maize (Hassan et al., 2023), made it an ideal system to evaluate SWIF-TE. The maize nested association mapping (NAM) founder lines, a set of 26 genome assemblies that represent the breadth of diversity in the species, were ideal for this benchmarking, as these genotypes have gold-standard genome assemblies (Hufford et al., 2021), pan-genome structural and homology TE annotation (Ou et al., 2024), available SV insertion data based on whole genome alignments that have been intersected with the TE structural and homology annotation (Munasinghe et al., 2023), and short read data from the same individuals used to generate the genome assemblies. Benchmarking was first done by comparing the inbred line Oh43 as the query genome and B73 as the reference genome. Oh43 has 11,830 insertions that are not present in the B73 reference genome assembly (Table 1) <u>(Munasinghe et al. 2023)</u>. The proportion of TIPs from different TE orders and major superfamilies mostly follows the genome-wide distribution, except for a higher proportion of *PIF-Harbinger* and *hAT* DNA transposons in the Oh43 TIPs than is observed in the genome-wide TE content.

**Table 1.** TE distribution in B73 and Oh43 and TIPs between Oh43 and B73 used as a gold standard TE data set for program evaluation.

| Order | Superfamily | B73 TE Count | Oh43 TE Count | Oh43 TIP Count Relative to B73 | B73 TE Proportion | Oh43 TE Proportion | Oh43 TIP Proportion Relative to B73 |
| --- | --- | --- | --- | --- | --- | --- | --- |
| TIR | DTA (hAT) | 77444 | 86552 | 3302 | 0.082 | 0.078 | 0.279 |
|  | DTC (CACTA) | 75732 | 95826 | 832 | 0.080 | 0.087 | 0.07 |
|  | DTH (PIF-Harbinger) | 89408 | 95360 | 3495 | 0.094 | 0.086 | 0.295 |
|  | DTM (Mutator) | 39421 | 44402 | 588 | 0.042 | 0.040 | 0.05 |
| LTR | RIL (LINE) | 7710 | 8256 | 194 | 0.008 | 0.007 | 0.016 |
|  | RLC (Copia) | 188391 | 210407 | 1203 | 0.199 | 0.190 | 0.102 |
|  | RLG (Ty3) | 329857 | 374907 | 1773 | 0.348 | 0.339 | 0.15 |
|  | RLX | 140282 | 189476 | 125 | 0.148 | 0.171 | 0.011 |
| **Total** |  | **948,245** | **1,105,186** | **11,830** |  |  |  |

Using the gold-standard set of insertions between Oh43 and B73, rates of true positive (TP), false negative (FN), and false positive (FP) insertions from SWIF-TE were calculated. True negatives (TN) were not calculated, as this would include any base pair in the genome that does not have a structural insertion (i.e. ∼2.3 billion base pairs in maize), which is not mathematically appropriate relative to the number of insertional polymorphisms. To get an idea of the true negative rate, we compared the SWIF-TE identified TIPs to the set of structural variant calls from the whole genome alignments between Oh43 and B73 that do not contain a TE sequence (termed noTESV in (Munasinghe et al., 2023)). These non-TE SVs should not be identified as TIPs by SWIF-TE. Our assumption here is that these positions with non-TE SVs are more likely to contain FP TIPs than other parts of the genome, so this comparison provides a conservative estimate of the TN rate. Out of the 11,557 noTESV calls, only 57 were called by SWIF-TE as FP, and the remaining 11,500 (99.5%) were TN.

Precision, sensitivity, FDR, and F1 scores were calculated from the TP, FN, and FP rates to benchmark SWIF-TE. Precision is the true discovery rate, while FDR is the false discovery rate. Sensitivity measures the proportion of identified insertions out of the total set of true insertions. Sensitivity is a result of both the evidence from the short read data (i.e., soft clipped reads at the insertion point) as well as the completeness of the TE library to determine that an insertion sequence is indeed a TE and not another inserted sequence. Finally, the F1 measure is the harmonic mean of precision and sensitivity. F1 is similar to accuracy but is useful because it does not require an estimate of TN, which was not quantified for this study. The initial testing was performed using short read data with 15x genome-wide coverage aligned in single-end mode. SWIF-TE successfully identified 1,438 TPs and 3,887 FPs between the Oh43 short reads and the B73 genome, resulting in a 27% precision rate and a sensitivity of 14%. Despite a relatively high FDR of 73%, the F1 score suggests a balanced trade-off between precision and recall.

The SWIF-TE program offers many options for optimizing performance. Optimal parameters will differ by species based on the complexity of the genome and the species’ TE landscape. Here, we report parameter testing in the context of the maize TE landscape. This type of testing should be done in any new species to reduce the overall number of FP and FN and increase the number of TP outputs from the program.

To begin parameter testing, different read inputs were tested, including single-end and paired-end reads. Paired-end read data has more information to uniquely align reads compared to single-end data, and potentially identify additional novel TE insertions (Alkan et al., 2011). To determine if alignment in paired-end mode improved the precision rate, we aligned the same Oh43 reads in both paired-end and single-end mode to the B73 reference genome assembly. Surprisingly, the single-end read data performed better than the paired-end data, with a 75% increase in sensitivity and only a 2.5% drop in precision using paired-end alignments (Figure 2a). Paired-end alignments require both reads to map, which can be difficult if one of the reads originates from within a TE not present in the reference genome assembly (i.e., a TIP). Further, if the sequenced fragment spans across the TE, the fragment size provided to the aligner is no longer accurate. This complexity could explain the higher precision using single-end alignments. Another alignment parameter that can impact the accuracy of SWIF-TE is the minimum alignment length to the genome and TE library. In some species, the dominating TE families are families with short terminal sequences (e.g., terminal inverted repeats (TIRs)), and in others, they are long (e.g., long terminal repeats (LTRs)). To determine the ideal alignment length for our evaluation dataset (i.e., maize 150bp reads), we tested alignment lengths from 30bp to 60bp, incrementing every 10bp. Across all of the different length values that were tested, 9% were FP in all cases, while 27% were FP at only one length. For maize 150bp reads aligned in single-end mode, the optimum alignment length was 40bp or more aligned to the genome and 40bp or more aligned to the TE library (Figure 2b). These parameters resulted in the lowest FDR (73%) and highest precision rate (27%). These parameters, however, also resulted in the lowest sensitivity rate (25%). Still, this was determined to be optimal as it has the best proportion of TP out of all of the total TIPs identified (TP / TP + FP). Extreme lengths (i.e., 20,40 and 60,40) caused a decrease in precision (0.053 and 0.081), most likely because, with 150bp reads, there was either too little sequence to accurately anchor into the TE library or genome at these lengths. The optimal alignment distance into the TE library and into the genome will depend on both the length of the raw reads as well as the species of interest.

**Figure 2.**
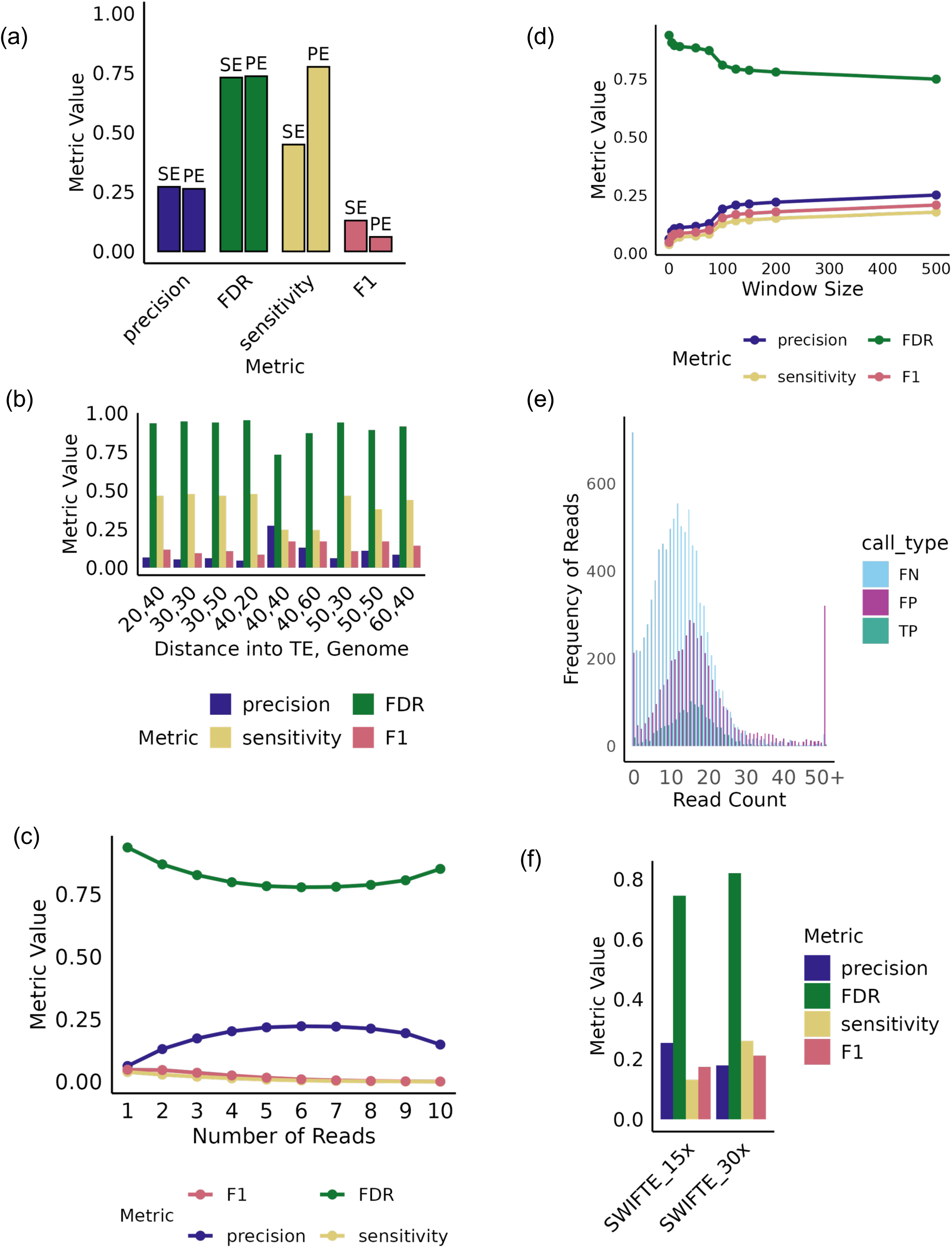
Impact of changing SWIF-TE parameters. For a) Percent change between single-end (SE) and paired-end (PE) mode for precision, sensitivity, FDR, and F1. b) Values for precision, sensitivity, FDR, and F1 for different distances into the TE library and the genome. c) Values of precision, sensitivity, FDR, and F1 for different numbers of reads supporting an insertion. d) Values for precision, sensitivity, FDR, and F1 for different window sizes to merge evidence. e) Frequency of split-read counts for FNs, FPs, and TPs. f) Values for precision, sensitivity, FDR, and F1 for both 15X and 30X genome-wide alignment coverage.

The number of reads required to support an insertion is another important parameter to optimize. In some cases, many reads supporting an insertion could come from a highly repetitive region resulting in a FP. Alternatively, a sequencing error could result in a FP when there is low read coverage. To determine the optimal number of supporting reads, we tested 1 to 10 reads of support from a 15x coverage dataset. Many of the FPs had very few supporting reads (1-2 reads; Figure 2c). Requiring at least 5 reads supporting the insertion was the most precise (0.27). At 6 or 7 reads of support, the precision was comparable to 5 reads of support, but the sensitivity was much lower. When requiring substantially more read support (i.e., 10 informative reads), sensitivity was quite low (0.000007), and precision was also lower than at 5 reads of support. In other datasets, the optimal number of reads to support an insertion will differ based primarily on the genome-wide coverage and will follow a linear relationship with coverage.

For each informative read, the position of the putative insertion supported by the read is provided. However, the exact base resolution of the insertion is not always correct. This can be due to allelic variation (i.e., SNPs and small InDels) that may result in truncation of the alignment to the genome or other alignment issues, which are common near TE insertion sites (Chu et al., 2020). Another adjustable parameter in SWIF-TE is the window size for collapsing informative reads to support a single insertion. Too large a window could collapse too many reads that support different insertions, while too small a window size might not accommodate the errors present in the data. We tested window sizes ranging from 0 to 500, increasing in 10-50bp increments (Figure 2d). A window of 100bp, which equates to approximately 50bp upstream and downstream of the actual TIP, effectively balanced high precision while maintaining low sensitivity (Figure 2d). This setting can be adjusted for different species, but it is crucial to remember that a window that is too large or too small can significantly affect both the sensitivity and precision.

The number of reads mapped to the region of interest where a TIP is called was also tested. Low mappability could indicate a region of the genome that is not mappable, whereas too high of coverage could indicate a highly repetitive region of the genome. The total number of reads mapped to the region in the genome was calculated across all TIP calls (Figure 2e). It was found that low mappability regions (0-1 reads mapped) and high mappability regions (35+ reads mapped) harbored a substantial amount of FN and FP, respectively. If cutoffs were implemented for these regions, 94% of TPs would be retained while 25% of FPs and 18% of FNs would be eliminated (Figure 2e).

Finally, we wanted to understand how sequencing depth affected SWIF-TE performance. We hypothesized that at higher whole-genome read coverage, SWIF-TE would identify more insertions because there would be more reads to identify insertions. We tested two different depths at 15x and 30x whole genome coverage. Surprisingly, SWIF-TE was more precise at 15x depth (27%), and precision dropped to 18% at 30x depth (Figure 2f). While precision dropped at the higher coverage, there was a substantial increase in sensitivity at 30x depth. Collectively, this shows that although more insertions overall were found at 30x coverage, there were more FPs present in the dataset (FDR increased from 75% to 82% at 15x vs 30x coverage). To balance the trade-offs between the different coverages and minimize the number of FPs moving forward, subsequent analyses were performed using the 15x coverage read dataset.

### SWIF-TE maximizes memory and runtime efficiency with minimal compromise on performance

Several other tools have been developed to identify TE insertions using short read data (Orozco-Arias et al., 2020; Sohrab et al., 2021; Yu et al., 2021). To benchmark the performance of SWIF-TE against existing programs, we looked for programs that used short read data as input and had the potential to scale to population analyses, as this was a major goal in the development of SWIF-TE. The programs that were benchmarked included TEfinder, TIP_finder, and TEMP2. TEfinder was designed for use in *Fusarium oxysporum*, a small genome, TIP_finder was designed for use in rice populations, and TEMP2 was designed for use in fly and human populations. To scale to large populations in species with large genomes, a program should have low runtime and memory requirements with minimal sacrifice on precision and sensitivity. Thus, benchmarking was performed based on runtime and memory usage to see which tools can effectively scale to population-level analyses. Benchmarking on precision, sensitivity, F1, and FDR was also used to evaluate the relative performance of the programs. For this benchmarking, the same 15x read data from Oh43 was run through each program and compared to the gold standard insertion set described above.

Runtime and memory usage are critical when performing population-level analyses, especially at the whole-genome level. Runtime is the amount of time a program takes to run and memory usage is how much RAM the program requires for the analysis. All of the programs were benchmarked on a high-performance Linux cluster with nodes that had 1995 GB of RAM and up to 128 threads available through the University of Minnesota Supercomputing Institute. SWIF-TE outperformed all other programs with regard to runtime and memory usage (Figure 3), using only 0.82 hours of runtime and 0.10 Gb of memory. The design of the algorithm, in which only 2Mb of alignments are processed at a time, makes it highly memory efficient compared to other programs that simultaneously load all data into memory.

**Figure 3.**
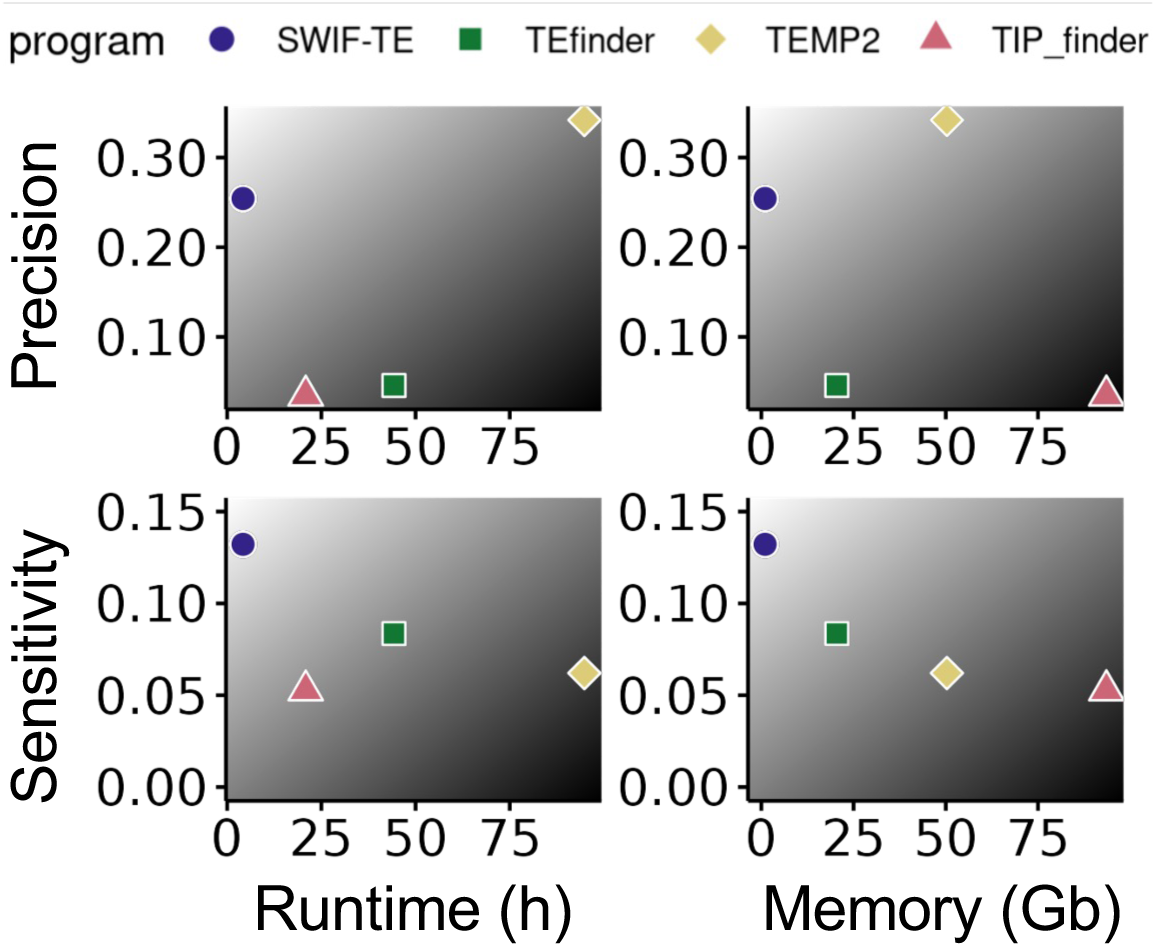
Benchmarking SWIF-TE and other available TIP calling software. Values for precision, sensitivity, runtime, and memory usage for SWIF-TE, TEfinder, TEMP2, and TIP_finder are displayed.

Performance metrics (i.e., TP, FP, FN) were also assessed using the gold standard TE data set described above for each of the programs. SWIF-TE identified the most TPs at 1,438 compared to 1,213 from TEfinder, 655 from TIP_finder, and 610 from TEMP2. As a result, SWIF-TE had the highest sensitivity at 0.13, followed by TEfinder at 0.08 (Figure 3). Precision is a function of both TP and FP. The most precise tool was TEMP2 at 0.32, with SWIF-TE following at 0.27 due to the higher rate of FP that is identified from SWIF-TE (Figure 3).

This benchmarking shows that SWIF-TE is an ideal program to use when the goal is to identify TIPs on a genome-wide scale, particularly in species with a high abundance of TEs and high variability in TE content. SWIF-TE maximizes memory efficiency and runtime with minimal compromise on precision. One program, TEMP2, outperformed SWIFT-TE in precision. However, this tool would take 41,400 hours and 25,000 Gb of memory to run in maize at the population level (i.e. 500 individuals) compared to SWIF-TE, which would only require 408 hours and 50 Gb of memory for the same size population.

### SWIF-TE identifies TIPs near genes with higher sensitivity and precision than TIPs further from the gene space

TEs can be found throughout the genome, including in genic and non-genic regions, and phenotypic variation has been associated with TIPs near or in genes (X. Li et al., 2024; Van’t Hof et al., 2016), but also with more distal TEs (Wang & Dooner, 2006). In maize, the majority of TIPs are located outside the gene space (Figure 4a), mirroring overall TE density in the genome (Hufford et al., 2021). However, with short read data, it can be difficult to obtain unique alignments in the highly repetitive non-genic portions of the genome. As such, we hypothesized that there would be bias in TIP identification based on proximity to genes. Indeed, polymorphic TEs inside or very near the gene space were more likely to be identified by SWIF-TE than those that are 5kb+ away from the gene (Figure 4a). Additionally, the ratio of TP to FP is greatest within genes and decreases moving away from the gene space (Figure 4b). These results combined demonstrate that, as expected, SWIF-TE not only has high sensitivity near or within genes but also has higher precision near or within genes as compared to the more repetitive non-genic space. This challenge with the alignment of short reads to repetitive genomes is a limitation of SWIF-TE and any other program that uses aligned short read data as input for TIP identification. Understanding these limitations of short read alignments is important to the biological interpretation of TIPs discovered with SWIF-TE, and likely any short read TIP-finding software.

**Figure 4:**
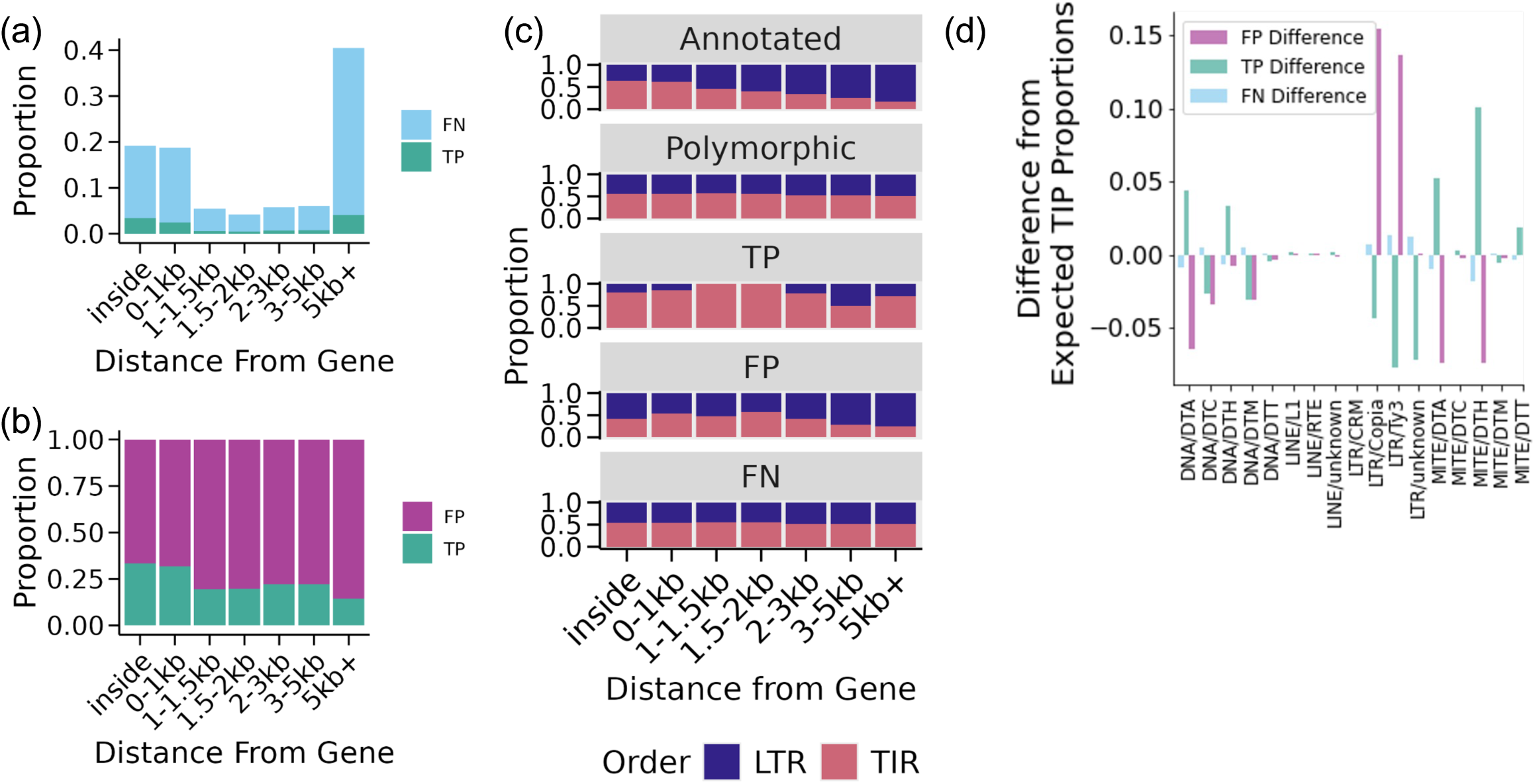
Influence of distance from gene and TE order on SWIF-TE TIP calls. a) Proportion of polymorphic TEs (FN & TP) in each distance from the gene space. b) Proportion of total TIPs called by SWIF-TE (FP & TP) and their distance from the gene space. c) Proportion of annotated, polymorphic, TP, FN, and FP LTRs and TIRs and their distance from the gene space. d) Differences from expected TIP proportions for FP, TP, and FN by superfamilies of TEs.

### SWIF-TE identifies TIRs with higher sensitivity and precision than LTRs

The ecosystems of where different TEs are located in the genome can vary across superfamilies and classes of TEs (Stitzer et al., 2021). For example, in maize, TIRs are more often located inside or near genes, while LTRs are more often located in the non-genic portion of the genome (Figure 4c). Given that there is a bias in sensitivity and precision (Figure 4a,b) based on proximity to the gene space, we sought to investigate the impact this would have on sensitivity and precision between TIRs and LTRs. To do this, the proportion of polymorphic TIR and LTR TEs from the gold standard set of TE polymorphisms was assessed. Surprisingly, among all the polymorphic TEs (i.e., TP and FN), the proportion of polymorphic LTRs and TIRs was relatively consistent from the genic to non-genic portions of the genome despite their variability in total abundance (Figure 4c). Within the TP TIP calls, TIRs were overall more abundant, except for 3-5 kb away from the gene space (Figure 4c), indicating that SWIF-TE has a higher sensitivity for TIRs than for LTRs regardless of proximity to the gene space. Furthermore, FP TIPs identified by SWIF-TE overall were more often LTRs than TIRs, especially as the distance from the gene space increased (Figure 4c). These results indicate the precision for TIRs is also higher than for LTRs.

Superfamilies within LTRs and TIRs have distinct features that could result in variability in their detection. Differences between the expected TIP superfamily frequency (TP and FN) and the actual TIP superfamily frequency (TP) could indicate how well SWIF-TE identifies certain superfamilies. Dramatic overrepresentation of FPs was observed for *Copia*, *Ty3*, and fewer than expected FPs were observed for miniature inverted repeat (MITE)/DTH and MITE/DTA, and the converse was observed for TP (Figure 4d). This overrepresentation was driven by a limited number of families, including DTHZM00389 and DTHZM00280 that were also frequently identified by PTEMD <u>(Kang et al. 2016)</u>, which also uses short reads to identify novel TIPs. This indicates that SWIF-TE has a more difficult time identifying *Copia* and *Ty3* elements, while it identifies more MITE/DTH and MITE/DTA elements correctly. *Copia* and *Ty3* elements are two of the most abundant families in the maize genome and are very repetitive, which can make mapping short reads to these sequences rather difficult, as compared to lower-abundance superfamilies.

### SWIF-TE captures a majority of the TIPs at the population level

A limitation of SWIF-TE is the high rate of FNs. In large populations of individuals, many of the non-reference TE insertions will be shared among individuals in the population (Figure 5a) (Anderson et al., 2019; Qiu et al., 2021). As such, the discovery of non-reference TE insertions using multiple individuals should reduce the overall amount of FNs discovered for a particular genotype. Unique TIPs that are only present in a single individual and were not initially discovered in that individual (Figure 5a) will not benefit from the population information.

**Figure 5.**
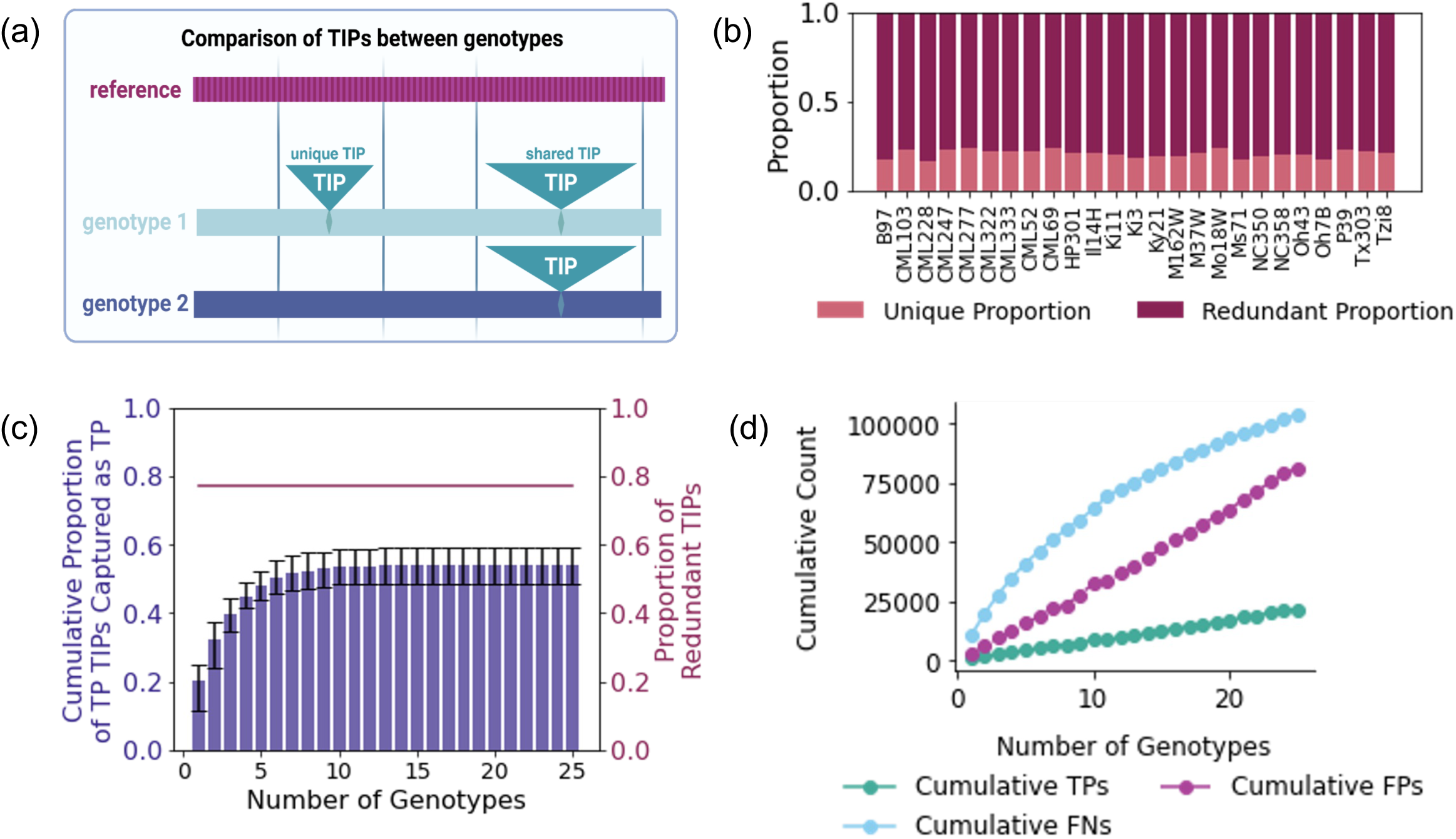
Population-level discovery of TIPs with SWIF-TE. a) Schematic of shared and unique TIPs between different genotypes. b) Proportion of unique and redundant TIPs in the 25 NAM lines compared to B73. c) Cumulative proportion of TIPs captured as true positive by SWIF-TE versus the number of genotypes added to the analysis. d) Cumulative counts of TP, FP, and FN versus the number of genotypes added to the analysis.

To evaluate the utility of such an approach, non-reference TEs in each of the 26 NAM genome assemblies were categorized as unique or shared (Figure 5a). On average, 73% of TIPs were shared with a TIP in at least one other line, and 27% were unique to each line (Figure 5b). Across all 26 NAM genome assemblies, 62,279 total of the gold-standard TIPs were unique to a particular line and, as such, have no opportunity for discovery in a different line, while 239,747 were shared in at least two lines. To begin looking at the benefits of using additional individuals to discover TIPs from a focal line, we first evaluate Oh43 as the focal line. Initially, only 22% of the TIPs that were identified from whole genome assembly were TP in the SWIF-TE output using only evidence from Oh43 short reads. With just one additional line, 13% more of the Oh43 TIPs were discovered (Supplemental Figure 1). After eight genotypes, a plateau was reached with 55% of the Oh43 TIPs discovered in the SWIF-TE output of at least one line (Supplemental Figure 1). A similar plateau was observed across all of the genotypes, with the number of discovered TIPs increasing from an average of 21% from self to 55% discovered across the population (Figure 5c). In adding additional genotypes, the cumulative number of TPs and FPs increases linearly, while the number of FNs is not linear and will reach a maximum (Figure 5d). This is because, with the addition of more individuals, SWIF-TE is able to capture more TPs that would have been FNs in a single line.

Across all of the NAM lines, there were 53,178 TIPs that were shared in at least two lines and were never discovered in any line using SWIF-TE. These recalcitrant TIPs were fairly evenly distributed across the genome, with 18% overlapping an annotated gene model, slightly higher than what is observed among all TIPs at 17%.

## DISCUSSION

Variation in the genome exists in the form of SNPs, small insertion/deletions (InDels), SVs, and TIPs. SNPs have historically been the most studied source of variation due to the abundance and the ease of identifying them computationally. SNPs can tag a lot of other sources of variation, and in doing so, represent that variation in efforts to dissect trait variation. However, not all TIPs are tagged by SNPs (Qiu et al., 2021). Directly identifying these variants will lead to improved understanding of the functional consequences of these variants (Qiu et al., 2021) with important implications for genome biology and the understanding of phenotypic variation (Domínguez et al., 2020; Zhang et al., 2025).

Previous tools for identifying TIPs were not designed for species with large genomes that have an abundance of variable TEs. Existing tools frequently suffer from long runtimes, high memory usage, and low precision, all of which are particularly problematic when attempting to scale to population-level analyses. In species with large genomes, the complexity increases substantially. SWIF-TE addresses these challenges by offering a powerful solution for TIP identification at the population level. Unlike existing tools, SWIF-TE is optimized for efficiency and scalability. By minimizing the trade-off between precision and performance, SWIF-TE enables the detection of TIPs across multiple genotypes while maintaining relatively high precision. Through its streamlined algorithm and memory-efficient design, SWIF-TE can process large genomic datasets quickly, making it an ideal tool for population-level analyses in species with large and TE-rich genomes. As demonstrated in our benchmarking, SWIF-TE outperforms other tools, providing both high precision and computational efficiency. This makes TIP identification in large, diverse populations feasible and accessible, offering new insights into how these genomic elements influence phenotypic traits and contribute to genetic diversity.

While SWIF-TE has been designed to improve computational performance, it is not without challenges. Storage of hundreds of SAM files for input into SWIF-TE can require substantial disk space, depending on the depth of sequencing coverage. The conversion of paired-end reads to single-end input can also require substantial computational time. SWIF-TE also has a high FN and FP rate, but some filtering for certain TE orders and using information across multiple individuals can improve this limitation. Parameter testing similar to what is reported here within new data sets and in different species using a gold-standard set of TEs is needed to optimize these parameters. Even with these challenges, SWIF-TE outperformed multiple other tools tested in runtime, memory usage, precision, and sensitivity. TIPs identified by SWIF-TE at the population level can then be scored using other more accurate methods <u>(Qiu</u> <u>et al. 2021; Gardner et al. 2017; Yu et al. 2021)</u> across all individuals in the population to recover missing information due to FNs. Overall, SWIF-TE is a powerful tool to use, and there are ways to reduce FP and FN.

## CONCLUSION

TE insertional polymorphisms have long been a challenging source of genomic variation to study. Here, we have described a newly developed software called SWIF-TE that enables efficient and precise identification of TIPs from short read data and can scale to population level analyses even in species with large genomes that have an abundance of variable TEs. Filtering parameters can be used to greatly improve the rate of FPs in the raw output, and information across individuals in the population is able to improve the discovery of novel insertions, reducing the FN rate. Further development of this software will focus on continuing to improve precision and sensitivity while maintaining the efficient memory and runtime that has been achieved with this software.

## METHODS

### Sequence Data

The maize reference genome (B73 version 5) was downloaded from MaizeGDB (https://download.maizegdb.org/Zm-B73-REFERENCE-NAM-5.0). Accessed 21 December 2021. The genome was indexed using bowtie2-build within bowtie2 version 2.3.4.1 (Langmead & Salzberg, 2012) with the –x option. The maize panEDTA TE library was downloaded from https://github.com/oushujun/PopTEvo/blob/main/TE_annotation/data/NAM.EDTA2.0.0.MTEC02052020.TElib.fa. Accessed 10 May 2023. Knobs, subtelomere repeats, and Helitrons were removed from the file before indexing using bowtie2-build within bowtie2 version 2.3.4.1 (Langmead & Salzberg, 2012) with the –x option. Short read sequence data used in the generation of 26 maize genome assemblies (Hufford et al., 2021) with 16-20X coverage were downloaded from the European Read Archive (ENA BioProject IDs PRJEB31061 and PRJEB32225). Accessed 20 January 2024.

### Sequence read processing and alignment

Short reads from each of the 26 genomes were trimmed using trimmomatic version 0.33 (Bolger et al., 2014) (PE –threads 16 –phred33 ILLUMINACLIP:all_illumina_adapters.fa:2:20:10 LEADING:3 TRAILING:3 MINLEN:25). Reads were aligned to the B73 reference genome assembly as both single-end reads and as paired-end reads. For single-end read alignment, the reads were first renamed to remove any information that linked paired reads to each other. Alignment to the B73v5 genome was completed using bowtie2 version 2.4.3.1 (Langmead and Salzberg 2012) with –p 24 –x –U –S – very-sensitive-local. For the paired-end alignment, trimmed reads were aligned to the indexed B73v5 genome using bowtie2 version 2.3.4.1 (Langmead and Salzberg 2012) with –p 24 –x –U –S –very-sensitive-local. Trimmed reads were also aligned to the indexed TE library using the same options for both single-end and paired-end alignments. Alignments were downsampled to either 15x or 30x genome coverage using samtools version 1.16.1 (H. Li et al., 2009) with the –s option, and the same reads that were sampled were included in analysis to the TE library. Before running SWIF-TE, picard MarkDuplicates (https://broadinstitute.github.io/picard/) was used to identify and remove duplicates in the SAM alignment file.

### Execution of SWIF-TE and other TIP calling programs

The downsampled 15x coverage Oh43 SAM file that was generated from single-end read alignment was used to initially run SWIF-TE and other TIP calling programs for performance benchmarking. For this benchmarking, SWIF-TE was executed using default parameters that included using the option 40,40 for minimum alignment length into the genome and minimum alignment length into the TE sequence.

Three other TIP calling programs were run for benchmarking relative to SWIF-TE, including: TEMP2 (Yu et al., 2021), TIP_finder (Orozco-Arias et al., 2020), and TEfinder (Sohrab et al., 2021). Each program was executed using the same 15x coverage Oh43 SAM input file that was generated from single-end alignments. Because each program reported differently on the insertion start and end, each program’s output was sorted by chromosome and then by the reported start position. If a range of positions was given for the insertion, the midpoint of the range was used as the insertion site position.

TEMP2 version 0.1.7 was executed within McClintock version 1.2.0 (Nelson et al., 2017) using default parameters. McClintock was downloaded from https://github.com/bergmanlab/mcclintock. Accessed 10 November 2022. The TEMP2 output was then filtered to contain only non-reference insertions for comparison to the other programs.

TIP_finder was cloned from https://github.com/simonorozcoarias/TIP_finder (Orozco-Arias et al., 2020). Accessed 26 January 2022. The TE fasta file was split into chunks of 0.338MB size, and TIPfinder was run in parallel with each chunk. TIP_finder output gives 10kb windows for the location of each insertion. Using the read alignment information, the middle position was identified for each of the TIP_finder insertional calls and used as the position of the TIP. TIP_finder was run using default parameters, using 16 threads in parallel on each B73 chromosome on each TE sequence separately before the output was concatenated together. Because TIP-finder was run in chunks, the runtime and memory usage were added together from each job submitted to the cluster for resource benchmarking.

TEfinder was cloned from https://github.com/VistaSohrab/TEfinder/tree/master (Sohrab et al., 2021). Accessed 21 December 2021. The TE library file was split into chunks of 0.338MB size and the reference genome was split into each chromosome. TEfinder was run using default parameters and on each TE sequence separately before the output was concatenated together. As with TIP-finder, runtime and memory usage were added together from each chunk for resource benchmarking.

Runtime and memory usage were calculated for each program using the seff (version 1.12) command in SLURM. Every program was run on the Agate Linux supercomputer at the University of Minnesota Supercomputing Institute, which has 1995 Gb of RAM on each node and 96 hours of runtime capacity. Each program was run on the max node on the UMN MSI Agate cluster. When using multiple processors, the runtime and memory were calculated per processor and then added together to get the final usage.

### Identification and Comparison to Gold Standard TE Insertional Polymorphisms Set

Evaluating the performance of SWIF-TE and other TIP calling programs required a gold standard set of TIPs for comparison. Previously, each of the NAM genome assemblies (Hufford et al., 2021) was aligned to the B73 reference genome assembly (Munasinghe et al., 2023) using the AnchorWave whole genome alignment program (Song et al., 2021), and all identified structural variants (SV) were classified relative to their TE content. For the gold standard TIP set, the SVs were filtered down to only those that were classified as TE=SV, where there is a 90% overlap between the SV and a single annotated TE. The TE=SV set was further filtered to include only those that were surrounded by alignable sequence and had unique AW Block IDs. Nested TE insertions were not retained in this filtering. Calls from each program that fell in unalignable sequence (from AnchorWave) were also removed.

A TP TIP called from the short read data was defined as one that is within a 100bp window around a TIP from the gold-standard set of TIPs. These overlaps were identified using the window program within bedtools version 2.29.2 (Quinlan & Hall, 2010), requiring a 100bp window using the –w 100 option. A FP TIP called from the short read data was defined as one that was not within a 100bp window of a TIP from the gold-standard set of TIPs using the same bedtools window. Finally, a FN was defined as a TIP in the gold-standard set that did not have a TIP called by the program within a 100bp window, and was identified using the window program within bedtools, and requiring no hits with the –v option. These classifications of TP, FP, and FN were used to calculate precision as TP/(TP+FP), sensitivity as TP/(TP+FN), F1 as (2*TP)/((2*TP)+FP+FN), and FDR as FP/(TP+FP).

### SWIF-TE parameter testing

There are a number of options within SWIF-TE that can be optimized based on the system and biological questions being addressed. These options were iteratively tested to determine the impact on rates of TP, FP, and FN TIP calls. The window size around the SWIF-TE identified insertion site was first tested to determine the appropriate window size for parameter testing. This was done using bedtools window –w –u version 2.29.2, outputting unique calls at incrementing steps between 0 and 500bp using the SWIF-TE output file and the true TE insertion dataset. It was determined that a window of 100bp around the SWIF-TE insertion site was appropriate for the data.

The first parameter within SWIF-TE that was tested was the number of reads of support required for a TIP call using a post-processing script to collapse reads within 100bp of each other and filter based on the number of reads supporting a TIP (see clean_SWIF-TE.sh on Github). Between 1 and 10 reads of support were required. Five reads of support was determined to be optimal and was used for all downstream parameter testing and population analysis.

Next, the impact of the average read depth was tested. The Oh43 BAM file from the single-end alignment was downsampled to 15x depth and 30x depth as described above. SWIF-TE was run on both 15x and 30x coverage read datasets using default parameters and requiring at least 5 reads of support. The results were benchmarked using the gold standard TIP dataset.

We also tested both paired and single-ended read alignments as input for SWIF-TE. The 15x depth Oh43 alignments from both paired-end alignments and single-ended alignments were input into SWIF-TE using default parameters, requiring at least 5 reads of support, and were again benchmarked using the gold standard TIP dataset.

Finally, the minimum alignment length to the genome and TE library was evaluated. The minimum distances of each of these were iterated at different levels, including: 20,40 (distance into TE, distance into genome), 30,30, 30,50, 40,20, 40,40, 40,60, 50,30, 50,50, and 60,40. These results were benchmarked using the gold standard TIP dataset.

### Genomic context of SWIF-TE calls

To test how distance from the gene space affected SWIF-TE precision, bedtools intersect (version 2.29.2) was used to intersect SWIF-TE calls at increasing intervals (0 to 5 kb and greater than 5kb) from B73v5 annotated genes (downloaded from https://download.maizegdb.org/Zm-B73-REFERENCE-NAM-5.0/archive/Zm00001e.1/Zm-B73-REFERENCE-NAM-5.0_Zm00001e.1.gff3.gz and filtered for ‘gene’). Accessed 15 May 2023.

To classify TIPs into their subsequent superfamilies and orders, annotated TEs from B73v5 were downloaded from https://download.maizegdb.org/Zm-B73-REFERENCE-NAM-5.0/Zm-B73-REFERENCE-NAM-5.0.TE.gff3.gz. Accessed 15 May 2023. These were used to create a dataframe to classify the TIPs identified by SWIF-TE into their superfamilies as only the family level information from the TE database is output from SWIF-TE.

## Data and Code Availability

All data used in this manuscript were from previous studies and are available at the links above. Code for running SWIF-TE and the analyses in this manuscript are available at https://github.com/HirschLabUMN/SWIFTE.

## DECLARATIONS

### Ethics approval and consent of participants

Not applicable.

### Consent for publication

Not applicable

### Availability of data and materials

All data generated or analysed during this study are included in this published article. The SWIF-TE and all analysis code are available at https://github.com/HirschLabUMN/SWIFTE.

### Competing interests

The authors declare that they have no competing interests

### Funding

This work was supported by the National Science Foundation (NSF) Grant No. IOS-1934384 and the Minnesota Agricultural Experiment Station (MAES). CCM was supported by the University of Minnesota DSI-MnDRIVE Graduate Assistantship.

## Author Contributions

CCM contributed to the data analysis, interpretation of the data, and wrote the manuscript. NSC, AEP, YQ, ER, and MM contributed to the data analysis and interpretation of the data. NMS and EBJ contributed to the design of the work and acquisition of funding. CNH contributed to the design of the work, acquisition of funding, and wrote the manuscript. All authors have read and approved the manuscript text.

## ABBREVIATIONS

FDR: false discovery rate
FN: false negative
FP: false positive
LTR: long terminal repeat
MITE: miniature inverted-repeat
NAM: nested association mapping
TIR: terminal inverted repeat
TE: transposable element
TIP: transposable element insertion polymorphism
TN: true negative
TP: true positive
SAM: sequence alignment map
SWIF-TE: short read whole genome insertion finder for transposable elements
SV: structural variant

## Acknowledgements

The authors acknowledge the Minnesota Supercomputing Institute (MSI) at the University of Minnesota for providing resources that contributed to the research results reported within this paper.

## FIGURE LEGENDS

**Supplemental Figure 1.**
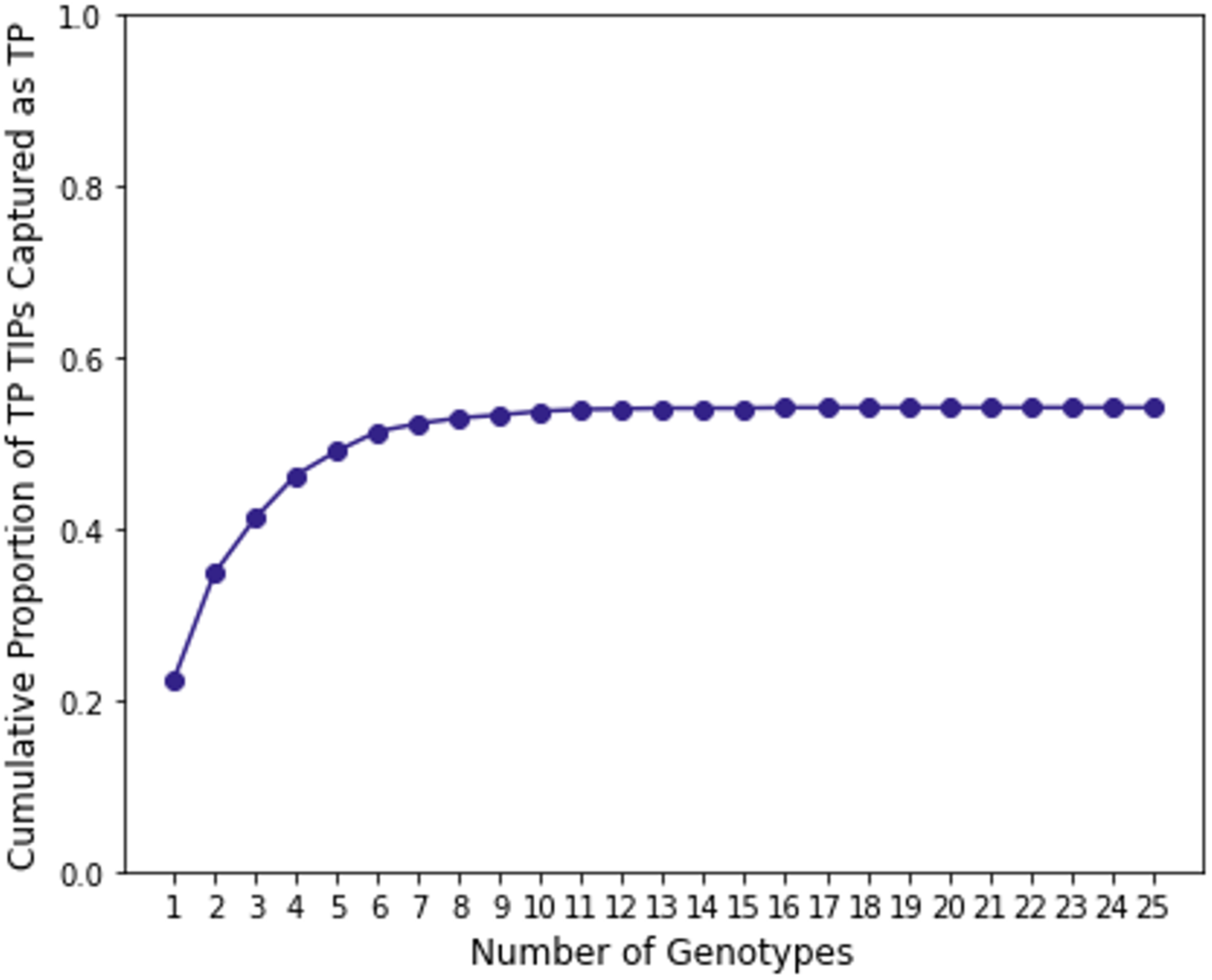
Cumulative proportion of TIPs captured by SWIF-TE in Oh43 versus the number of genotypes added to the analysis.

## Notes

### Competing Interest Statement

The authors have declared no competing interest.

